# Going Beyond the Point Neuron: Active Dendrites and Sparse Representations for Continual Learning

**DOI:** 10.1101/2021.10.25.465651

**Authors:** Karan Grewal, Jeremy Forest, Benjamin P. Cohen, Subutai Ahmad

## Abstract

Biological neurons integrate their inputs on dendrites using a diverse range of non-linear functions. However the majority of artificial neural networks (ANNs) ignore biological neurons’ structural complexity and instead use simplified point neurons. Can dendritic properties add value to ANNs? In this paper we investigate this question in the context of continual learning, an area where ANNs suffer from *catastrophic forgetting* (i.e., ANNs are unable to learn new information without erasing what they previously learned). We propose that dendritic properties can help neurons learn context-specific patterns and invoke highly sparse context-specific subnetworks. Within a continual learning scenario, these task-specific subnetworks interfere minimally with each other and, as a result, the network remembers previous tasks significantly better than standard ANNs. We then show that by combining dendritic networks with Synaptic Intelligence (a biologically motivated method for complex weights) we can achieve significant resilience to catastrophic forgetting, more than either technique can achieve on its own. Our neuron model is directly inspired by the biophysics of sustained depolarization following dendritic NMDA spikes. Our research sheds light on how biological properties of neurons can be used to solve scenarios that are typically impossible for traditional ANNs to solve.

## 1 Introduction

The *point neuron* model has been the primary neuron model used in computational and theoretical studies of neurons for over 100 years. Originally proposed by Louis Lapicque in 1907 (Lapique, 1907) the model assumes a simple linear integrate and fire mechanism. This continuous time mechanism formed the basis for Rosenblatt’s discrete Perceptron model (Rosenblatt, 1958), and continues to form the basis for current deep learning systems and artificial neural networks (ANNs) (McClelland et al., 1986, LeCun et al., 2015). In standard ANNs each neuron computes a linear weighted sum of its inputs, followed by a non-linearity (Figure 1 left). In contrast, *pyramidal neurons*, which comprise most cells in the neocortex, are significantly more sophisticated than point neurons, and demonstrate a wide range of complex non-linear dendrite-specific integrative properties (Spruston, 2008) (Figure 1, right). In this paper we focus on active dendritic properties where different dendritic branches on a single neuron act as independent pattern detectors (Antic et al., 2010, Major et al., 2013).

**Figure 1:**
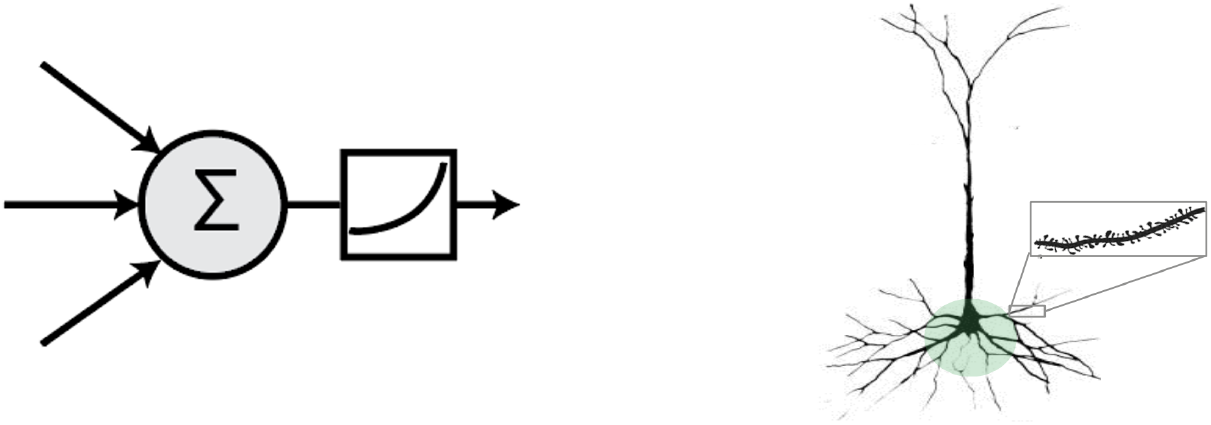
**Left:** the point neuron prevalent in most ANNs today simply computes a linear weighted sum of its inputs followed by a non-linearity. **Right:** a pyramidal neuron in the brain exhibits a vastly more complex structure.

Computational models have suggested that these *active dendrites* can be incorporated into neuron models by treating each dendritic neuron as containing multiple layers of point neurons (Poirazi and Papoutsi, 2020, Beniaguev et al., 2021). Although these studies show that such neurons have greater complexity, computational power and capacity than single point neurons, full networks constructed from these dendritic neurons are not fundamentally more powerful or capable than large multi-layer networks constructed from point neurons. Because of this fact most ANNs continue to use the point neuron model.

In this paper we show that dendrites impart fundamental computational benefits beyond a simple increase in complexity or computational power. We incorporate several properties of biological neural networks into an ANN: active dendrites, local inhibition and sparsity, and complex synapses. We then explore the impact of each of these changes in *continual learning* scenarios. In these scenarios the network is trained on a continuous sequence of tasks (McCloskey and Cohen, 1989, van de Ven and Tolias, 2019). Standard ANNs perform extremely well in batch training but do not perform well in continual learning. In particular they are unable to learn a new task without erasing what they previously learned, a phenomenon known as *catastrophic forgetting* (McCloskey and Cohen, 1989, French, 1999, Parisi et al., 2019). Continual learning is a core capability, central to intelligent systems, and something that humans perform naturally.

Our work builds on the initial HTM neuron model used in Hawkins and Ahmad (2016), which in turn models the depolarization effects of active dendrites (Antic et al., 2010, Major et al., 2013). The HTM neuron contains two different dendritic zones and receives two sources of input: a bottom-up feedforward input and a second context input. The sparse contextual input acting on a neuron acts as a prediction which biases the neuron and makes it more likely to become active.

Our primary goal in this study is to show a proof-of-concept working system that illustrates how these active dendritic concepts can be incorporated into a deep learning system. The main contributions are as follows:

- We summarize some of the existing experimental and theoretical work on active dendrites, sparse representations, and continual learning.
- We propose a new deep learning architecture that incorporates dendrites and sparse representations.
- We show experimental results on a standard continual learning benchmark, permutedMNIST. The results show that networks with active dendrites can retain a significant fraction of past information.
- We analyze the results and suggest reasons why active dendrites and sparse representations help with catastrophic forgetting.

## 2 Background

### 2.1 Active Dendrites

A prototypical pyramidal neuron has an extensive dendritic arbor containing thousands of synapses, each receiving input from other neurons (Bentivoglio and Swanson (2001), Ziehen (1895), Kandel (2012)). The point neuron model—used in most current deep learning networks—says that all these synapses have a linear impact on the cell. The cell would fire if its total inputs exceed a threshold and then resets. However the vast majority of the synapses are distal (far from the cell body) and individually have minimal impact on the cell. Only proximal synapses (close to the cell body) have a linear impact on the neuron. Distal dendritic segments instead process groups of synapses locally in a non-linear fashion, and are known as active dendrites (Major et al., 2013). Empirical evidence (Larkum et al. (1999), London and Häusser (2005), Branco and Häusser (2010), Antic et al. (2010), Major et al. (2013)) suggests that each distal dendritic segment acts as a separate active subunit performing its own local computation. When the number of active synapses within a small local region reaches above a threshold, the segment initiates a dendritic spike, independent of activity elsewhere.

Some modeling studies propose that active dendritic segments act as separate layers (Poirazi et al. (2003), Poirazi and Papoutsi (2020), Jadi et al. (2014), Beniaguev et al. (2021)) and suggest that each pyramidal neuron is equivalent to a multi-layer network of point neurons. Theoretical analysis of these models show that individual neurons with active dendrites have greater capacity than point neurons (Poirazi et al., 2003). Although this suggests that neurons are significantly more powerful than originally thought, from a deep learning point of view, the model does not provide a compelling justification to incorporate active dendrites. It is sufficient to simply implement a deeper ANN composed of point neurons.

In our model we exploit an additional property of dendritic spikes. Dendritic spikes travel to the cell body but are typically insufficient on their own to cause the cell to fire. Instead, a dendritic spike can depolarize the neuron for an extended period of time, sometimes as long as half a second (Antic et al., 2010, Major et al., 2013, Gao et al., 2021). During this time, the cell is significantly closer to its firing threshold and any feedforward input is more likely to make the cell fire. This suggests that active dendrites have an indirect *modulatory* impact on the cell’s response, with a very different role than feedforward inputs. The sustained depolarized state has been called a “predicted state” (Hawkins and Ahmad, 2016) or a “prepared state” (Antic et al., 2018). Active dendritic segments typically receive contextual input that is a different input than received in proximal segments. These context signals can arrive from other neurons in the same layer, neurons in other layers, or in the form of top-down feedback. Recent experimental evidence has shown that the input on active segments can drive context dependent activity (Takahashi et al., 2020). In the HTM model (Hawkins and Ahmad, 2016) we showed that contextual input on active dendrites can invoke context-specific activity and lead to a powerful predictive sequence memory. In this paper we discuss how these notions can be extended to context-dependent activity in deep learning networks.

### 2.2 Sparse Representations

Biological circuits and neocortical neurons exhibit sparsity in terms of both their 1) activations and 2) connectivity. In regards to activation sparsity, previous studies showed that in the neocortex relatively few neurons spike in response to a sensory stimulus, and that this is consistent across multiple sensory modalities, i.e., somatosensory, olfactory, visual and auditory (Attwell and Laughlin, 2001, Barth and Poulet, 2012, Liang et al., 2019). While the mechanisms maintaining sparsity, and the exact sparsity at the level of the individual neuron, remain to be answered fully, sparsity is a well-documented cortical phenomenon. Sparsity is also present in neural connections: cortical pyramidal neurons show sparse connectivity to each others and receive only few excitatory inputs from most surrounding neurons (Holmgren et al., 2003).

In computational modeling, sparse neural activity in the brain is translated into sparse representations: vectors where most of the entries are off (i.e., equal to zero) (Olshausen and Field, 1997, Cui et al., 2017). Just as in dense representations, individual entries can correspond to the presence of certain features, such as an edge in a particular position in an input image, hence sparse representations encode semantics. One advantage of sparsity in representations is that vectors for two separate entities have low *overlap*, which means the set of features/entries that are non-zero in both vectors is small. Previous studies with ANNs have found that sparse representations lead to more noise robustness than do dense representations, and slight perturbations in the input are less likely to hinder a trained pattern recognizer (Ahmad and Hawkins, 2016, Ahmad and Scheinkman, 2019). The idea of low representation overlap among unrelated inputs is particularly useful when learning in sequence. If the representations of two inputs from different “tasks” have near-zero overlap, it’s easier for the learner to recognize which task a given input most likely corresponds to, by simply responding to which entries are non-zero, and can make a prediction accordingly.

### 2.3 Continual Learning

Continual learning is the ability to acquire new knowledge over time. In contrast with biological networks, typical deep learning networks perform poorly in this setting. We review previous work in continual learning from two main categories: 1) regularization-based and 2) subnetwork-based approaches. The former regulates plasticity levels throughout the network during the course of training. In recent years, two of the most prominent examples of regularization are Elastic Weight Consolidation (EWC) (Kirkpatrick et al., 2017) and Synaptic Intelligence (SI) (Zenke et al., 2017). Both approaches (EWC and SI) estimate how relevant each parameter of the network is towards solving each previously encountered task. Motivated by the complex synapse structures seen in biology, SI uses two parameters per weight with internal dynamics that depend on the relevance of each weight to each task.

Subnetwork-based approaches are concerned with identifying subpopulations of neurons that each learn one of the many tasks in the sequence. Gated Linear Networks (Veness et al., 2021) and Dendritic Gated Networks (Sezener et al., 2021) are examples of this type of approach and work by applying a gating mechanism that selects subnetworks based on the input. In another example, Wortsman et al. (2020) present Supermasks, in which each task is designated a subset of neurons in the network. While Supermasks can learn hundreds of Omniglot (Lake et al., 2015) classes in sequence, computing the appropriate subnetwork is largely decoupled from the inference procedure and must be done on the side, a limitation of this approach. Context-dependent Gating (XdG) (Masse et al. (2018)) selects predetermined subnetworks of neurons, but exact task information must be provided during training and testing. Meta-learning can also be a useful tool to generate subnetworks: both Javed and White (2019) and Beaulieu et al. (2020) employ meta-learning to learn continually, and sparse subnetworks tend to emerge without explicit hard-coding. In Beaulieu et al. (2020) the systems learn a modulation network that selects subpopulations of neurons, an algorithm referred to as ANML. One limitation of this approach is that the only weights that change during continual learning are the output weights; the vast majority of the networks stays fixed.

## 3 Model

In this section we describe our method for translating the above biological properties into a deep learning architecture. We can summarize some of the key differences between pyramidal neurons and the point neuron model as follows:

1. Pyramidal neurons integrate inputs at multiple dendritic segments, whereas a point neuron has a single integration zone.
2. Proximal and distal dendritic inputs have different impacts on a neuron’s voltage and output, but in a point neuron, all inputs are treated like proximal inputs. Distal inputs on active dendrites can modulate a neuron’s response, making it more likely to fire.
3. Proximal and distal inputs may come from separate sources, whereas the point neuron assumes there is a single source of synaptic inputs.
4. Neural activity is generally sparse, whereas the activity is dense in most standard ANNs.

In our model, we strive to narrow these differences in the context of a deep learning system. In the rest of this section we describe our modified architecture where neural activity is dependent on both proximal (feedforward) and distal (context) inputs. The system is differentiable such that the entire network can be trained end-to-end using standard backpropagation. We later test our hypothesis that these context-dependent responses are useful for continual learning. Our implementation is available online^1^.

### 3.1 Active Dendrites Neuron

To turn these ideas into a formal computational model, we present the *Active Dendrites Neuron* which goes beyond the point neuron in how it implements neural computation. Our Active Dendrites Neuron (Figure 2) receives two sources of input, analogous to the proximal and distal inputs in pyramidal neurons. The feedforward inputs are processed just as in a point neuron, by computing a linear weighted sum of inputs. In addition, a set of dendritic segments on each neuron process a context (distal) input, and the subsequent dendritic output modulates the neuron’s response. The end result is a neuron where the magnitude of the response to a given stimulus is highly context-dependent.

**Figure 2:**
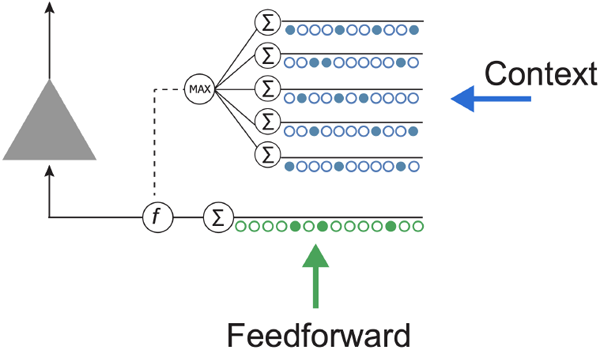
Illustration of a single Active Dendrites Neuron. Feedforward weights (green) receive regular feedforward input while dendritic segments (blue) receive a context vector. A single activation is further chosen after all dendritic segments compute an activation value, which thus modifies the linear weighted sum computed by feedforward weights.

Given a feedforward input vector ***x***, our neuron computes 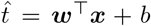 where ***w*** and *b* are the feedforward weights and bias of the neuron, respectively. Each dendritic segment *j* computes a linear weighted sum of a context vector 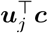, where ***u***_*j*_ are the weights for that segment. We select the segment with the strongest response to the context vector and compute the *dendritic activation, d*, used to modulate the neuron: 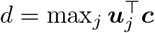 (The method we use to compute the context vector, ***c***, is described in Section 3.3.)

The output of a single Active Dendrites Neuron is:

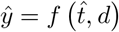

where *f* is a *modulation function* that modifies 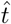 by the dendritic activation. In this paper we choose *f* to perform sigmoidal gating: 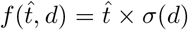, where *σ*(·) is the sigmoid function which takes a real number and squishes it into the range [0, 1]. Thus:

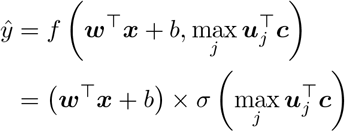

Here, a strong positive dendrite response to the context vector will retain the feedforward activation since the sigmoid will be close to 1. Weak or negative responses to the context vector will significantly reduce the activation since the sigmoid will be significantly lower than 1. There are many variations of the above formulation that are possible. We found that the network worked best when we selected the dendrite activation with the largest absolute value and retained the sign in *d*. This allows a strong negative response to more easily turn the neuron off (see Appendix A for more details).

### 3.2 Sparse Activations

We apply a *k*-Winner-Take-All (*k*WTA) function (Ahmad and Scheinkman, 2019) as our choice of non-linear activation in each hidden layer:

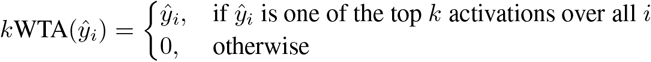

where *i* indexes neurons in the same layer.

The effect of *k*WTA is to pick out the top *k* activation values and drop all others to zero. The dendritic segments, by modulating the feedforward output, have a large impact on which neurons actually win. The *k*WTA layer thus results in sparse activity patterns that are highly context dependent. Note that in (Ahmad and Scheinkman, 2019) we also tested the impact of sparse weights. In this paper we found that 50% weight sparsity worked well, but we have not yet performed an exhaustive hyperparameter search to optimize weight sparsity.

### 3.3 Computing the Context Vector

In our model neural activity is modulated based on how well their dendritic segments detect a particular context vector, and the subsequent *k*WTA activation preferentially selects the up-modulated neurons. In a continual learning scenario, the context vector can serve as a task identifier so that neurons can be modulated for each task.

In order to apply our network to a continual learning dataset we need to compute an appropriate context vector. There is a large space of possible context vectors that could be used, and the exact choice can significantly impact overall network accuracy. In this paper we chose to use a simple prototype method to infer the context vector (Rosch, 1975, Snell et al., 2017). In this scheme a single vector is chosen to represent each task (see Figure 3). We implemented two different variations of the prototype method.

**Figure 3:**
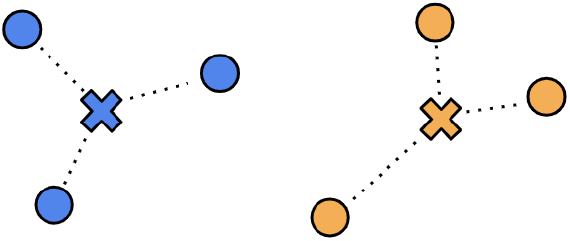
An illustration of the prototype method for computing context vectors. The blue circles are training samples in input space for task A, while the orange circles are training samples for task B. The blue cross is a vector that represents the prototype for task A, and the orange cross represents the prototype for task B. At test time, we identify the closest prototype to each input vector and use it as the context vector.

#### Training method 1 (task information provided)

In the first method we assume that the system receives task information during training. During training, all training samples for a particular task are assigned a single prototype context vector. We compute the prototype vector for task *t* by taking the element-wise mean over all the training samples across all features:

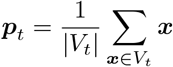

where *V*_*t*_ gives the set of all data samples ***x*** that the model observes to train on task *t*. The dimensionality of the context vector is thus identical to the dimensionality of the input vectors. This context vector is specific to each task and agnostic to the target label.

#### Training method 2 (task information not provided)

In the second method we relax the constraint that the identity of the task is given during training and implement prototypes that are automatically inferred during training in an online manner. To achieve this we use a statistical clustering approach that builds context prototypes on the fly. When the system receives a new batch of training samples from a task, we use an unpaired multivariate *t*-test to compare the current samples to previously-observed training samples. If the new batch of samples is similar to earlier training samples, they are assigned to an existing prototype. If not, the new batch of samples is assumed to be a new task, and a novel prototype is instantiated. In this case, there isn’t necessarily a one-to-one mapping between tasks and prototype context vectors. More details on this method are described in Appendix B.

#### Testing (selecting prototypes during testing)

For both of the above methods, at test time we do not provide any task information to the system. Instead it must dynamically choose the correct context vector and provide that to the network. We do this by selecting the closest prototype vector to each test example (using Euclidean distance) as the context vector. That is, for a test example ***x***^*′*^, the chosen prototype is:

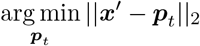

computed over all prototypes ***p***_*t*_ stored in memory.

### 3.4 Active Dendrites Network Architecture

Figure 4 shows the Active Dendrites Network that we used for our continual learning experiments. In each hidden layer, all neurons are Active Dendrites Neurons, and the network is trained end-to-end with standard backpropagation. We make two notes: first, only the neurons that were selected by the *k*WTA function will have non-zero activations (and thus non-zero gradients). Therefore during the backward pass only the weights corresponding to those winning neurons will be updated. Second, for each of those winner neurons, only the dendritic segment *j* that was chosen by the max operator is updated; all other segments ***u***_*j′*_ for *j*^*′*^ *≠ j* remain untouched. Thus a very small sparse subset of the full network is actually updated for each input. We hypothesize that a functional specialization will emerge where different dendritic segments will each learn to identify specific context vectors. This in turn will reduce interference during training between different tasks and thereby limit catastrophic forgetting. Indeed, since most dendritic segments that don’t respond to a specific context will not be updated, any context-dependent modulation of the neuron should be preserved from task to task. This aspect makes our Active Dendrites Network a suitable contender for learning continually.

**Figure 4:**
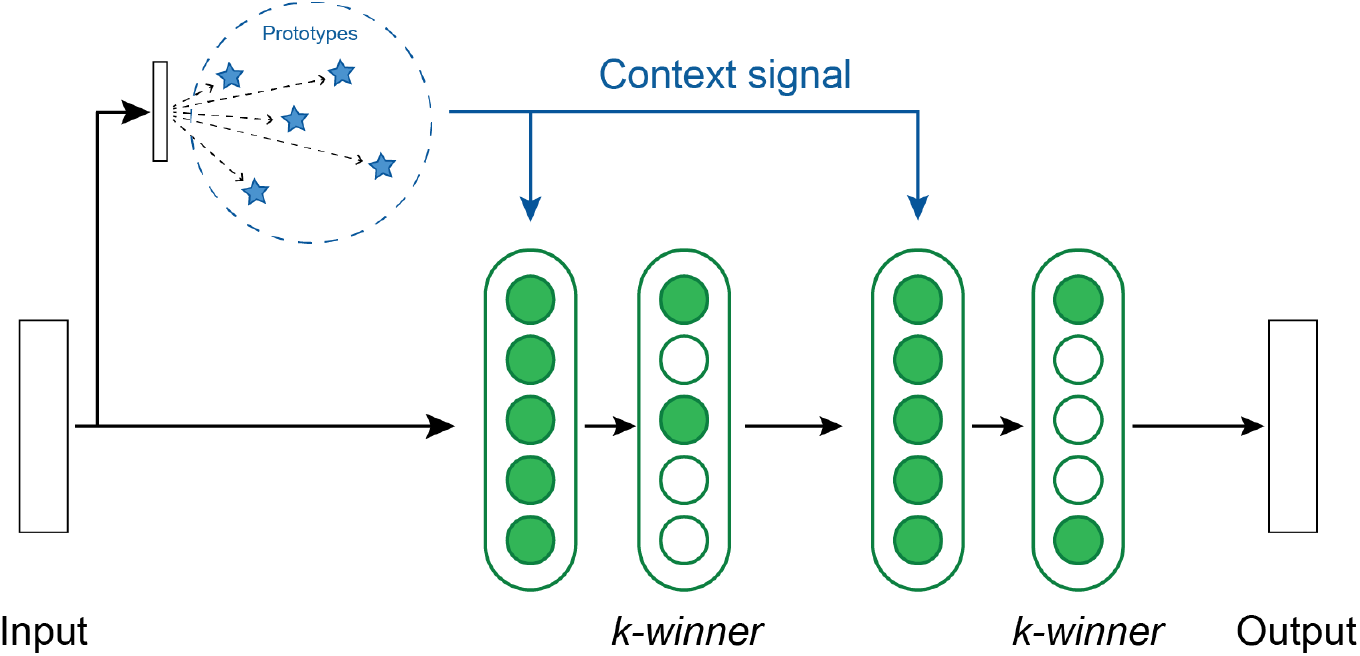
An overview of the base network structure used in our experiments. There are two layers of hidden units, each with a k-winner-take-all activation function. A context vector is computed from each input by locating the nearest prototype vector. The dendrites in each layer receive this context vector as input.

## 4 Results

A typical scenario in continual learning consists of training an ANN on a number of discrete tasks in sequence. Once the network has been trained on a particular task, it does not encounter that task again. The goal is to learn the tasks in sequence without forgetting previously-learned tasks.

We apply our Active Dendrites Network to permutedMNIST (Goodfellow et al., 2013), a common benchmark in continual learning, and discuss our findings in this section. In permutedMNIST, each task requires classifying images of handwritten digits from 0–9 just as in regular MNIST, except each task also applies a unique pixel-wise permutation to all images while maintaining the categories for each image. Consequently, the data distribution of each task changes, and ANNs are generally not permutation-invariant and thus forgetting occurs. Since the MNIST dataset contains 50, 000 training images, there are 50, 000 training images for each task. When trained on *T* consecutive tasks, the network is trained on a total of *T ×* 50, 000 images. Once training has completed, the network accuracy is calculated using a test set consisting of all *T* permutations applied to the MNIST test dataset.

### 4.1 Results on permutedMNIST

We used the network structure shown in Figure 4. Our network is composed of two hidden layers with 2,048 Active Dendrites Neurons each plus a final output layer with 10 neurons. We chose these network layer sizes to be similar to previous studies that report results on this dataset (Kirkpatrick et al., 2017, Zenke et al., 2017, Masse et al., 2018).

We train our model to learn up to 100 tasks in sequence. The network is tested at the end of training by computing accuracy on the test set for all tasks. When attempting to learn *T* consecutive tasks, the hidden neurons are equipped with *T* dendritic segments each to give it sufficient capacity to recognize a unique context vector for each task. We report accuracy numbers by averaging over 8 independent runs each with a randomly-picked seed. (Appendix D contains the hyperparameters used for each experiment.)

As shown in Figure 5 (left), we achieve accuracies of 94.6% and 81.4% on 10 and 100 consecutive permutedMNIST tasks, respectively, when context is provided during training, and accuracies of 94.3% and 76.9% when context needs to be dynamically inferred during training. Since there are always 10 categories, chance accuracy is 10% independent of the number of tasks. This demonstrates that the network is able to retain the majority of the knowledge from previous tasks. Note that a standard feedforward network performs poorly on this benchmark (Kirkpatrick et al., 2017, Zenke et al., 2017, van de Ven and Tolias, 2019) (see also Section 4.3 for more direct comparisons).

**Figure 5:**
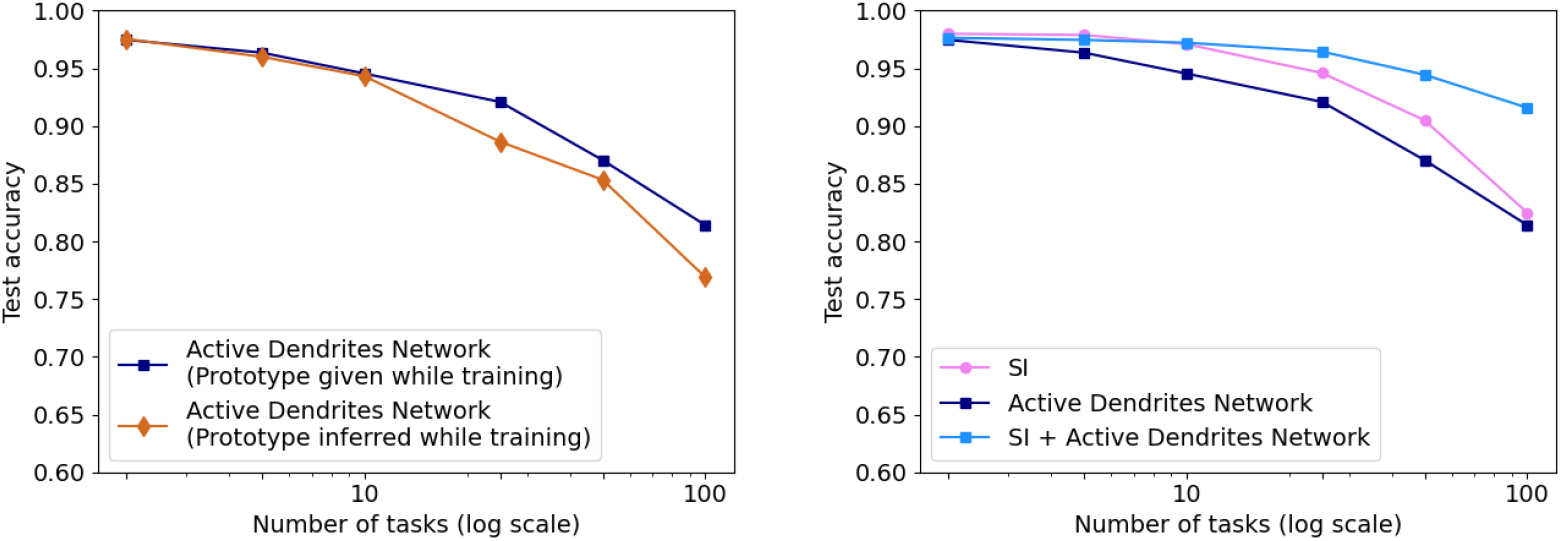
**Left:** The accuracy of our Active Dendrites Networks when learning 2, 5, 10, 25, 50, and 100 permutedMNIST tasks in sequence. We show results using both prototype methods while training: when the the model is provided with a prototype, and when it must infer the vector in an online manner. **Right:** The accuracy of the Active Dendrites Network and SI. The accuracy when combining SI + active dendrites is greater than either one on its own.

We also compare the results with SI (Zenke et al., 2017) (see Section 2.3). SI is motivated by the complex structure of biological synapses and known to do well on this benchmark. SI operates solely at the level of synapses: it maintains an additional parameter per weight that controls how fast that weight adapts to specific tasks. Since our architecture operates at the neuron and network levels, the two approaches are complementary techniques and can be combined. Figure 5 (right) shows the benefits of this combination. The accuracy of Active Dendrites Networks combined with SI improves to 97.2% and 91.6% accuracy on 10 and 100 consecutive tasks, respectively. Combining the two leads to higher accuracy than either method on its own. This suggests that biological mechanisms at the synapse, neuron, and network levels can operate together to handle continual learning. Note that SI as described in Zenke et al. (2017) requires knowledge of the task during training, as such we only combine it with our first prototype method. It may be possible to remove this restriction, and is a direction for future research.

### 4.2 Are Dendrites Invoking Subnetworks?

Our hypothesis behind incorporating active dendrites and *k*WTA is for the former to up- and down-modulate individual neuron’s activations, and for the latter to use this modulation to activate subnet-works that correspond to each task. We tested this hypothesis by analyzing the representations of each layer of Active Dendrites Neurons for a network trained on 10 tasks. Figure 6 shows the average activation frequency per task for the first 64 neurons in the second hidden layer after applying *k*WTA. Looking horizontally across the rows, each task appears to select a different sparse subset of neurons. Looking vertically, each neuron appears to activate frequently only for a small fraction of tasks.

**Figure 6:**
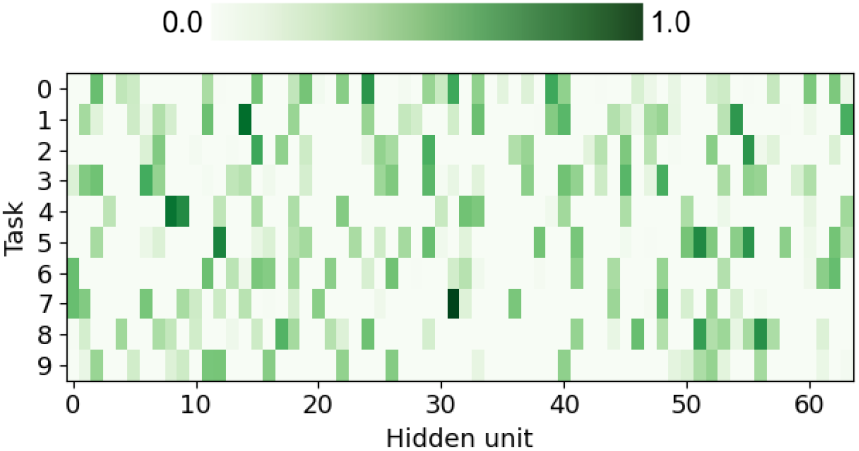
The fraction of instances for which each of the first 64 hidden units in the final hidden layer became active (after applying *k*WTA), when training an Active Dendrites Network on 10 permutedMNIST tasks. This figure separates instances by task, and uses 5,000 randomly-chosen test examples across all tasks. Note that each hidden layer contains 2,048 hidden units, but we show just 64 for ease of visualization.

To quantify this further we computed the mean cosine similarity of sparse activations for the entire layer, across all pairwise tasks. A low cosine similarity between any two tasks implies that the representations for the tasks are different and that the corresponding sub-networks are also different. Figure 7 shows the matrix of cosine similarities plus the average within-task and across-task similarities. According to this measure, it appears that the network has indeed learned to invoke non-overlapping subnetworks for different tasks.

**Figure 7:**
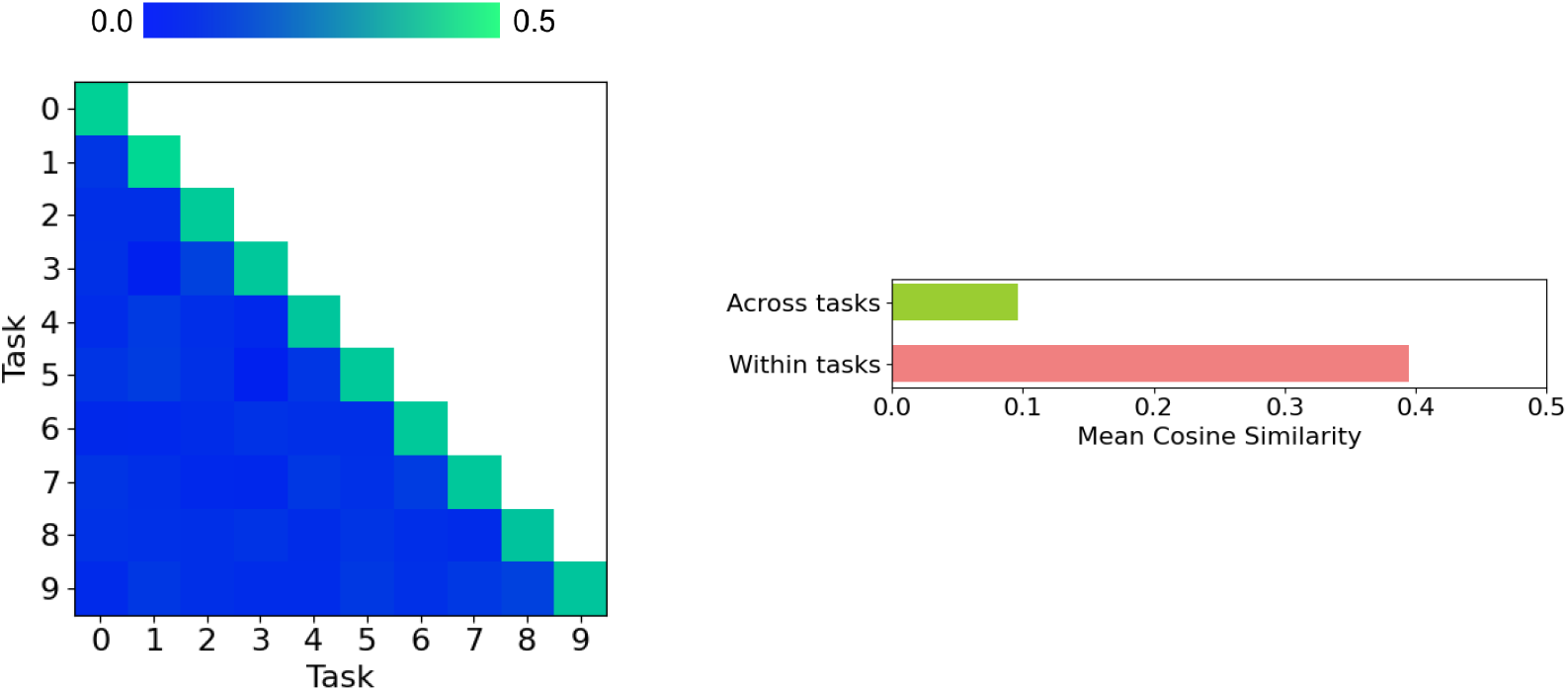
Mean pairwise cosine similarity values of hidden representations in the penultimate layer of an Active Dendrites Network after training on 10 permutedMNIST tasks in sequence. This figure uses 5,000 randomly-chosen test examples across all tasks. **Left:** Pairwise comparison between tasks where cell *i, j* gives the mean pairwise cosine similarity values across all pairs from tasks *i* and *j*. **Right:** Average across- and within-task cosine similarities (error bars omitted).

Even though subnetworks of neurons become active, what is the effect of the dendrites of a single neuron? In Figure 8, we zoom in on a few Active Dendrites Neurons and their responses (i.e., dendrite activations) to different context vectors before and after learning 10 permutedMNIST tasks in sequence. At the beginning of training the responses are random, with scattered positive, negative, and near-zero responses. After training, most responses are weak and only a few are very positive. We posit that with the *k*WTA layer selecting only the strongest neurons, dendrites don’t have much incentive to down-modulate the neuron to induce near-zero activation value. The *k*WTA effectively nullifies the activations of non-winners. The dendrites primarily need to up-modulate the neuron when it needs to become active as part of a subnetwork. This is consistent with the depolarization effect of dendritic spikes, which only up-modulate neurons. Across the neurons we have shown, dendrites only have strong responses to a few contexts as different neurons participate in different subnetworks.

**Figure 8:**
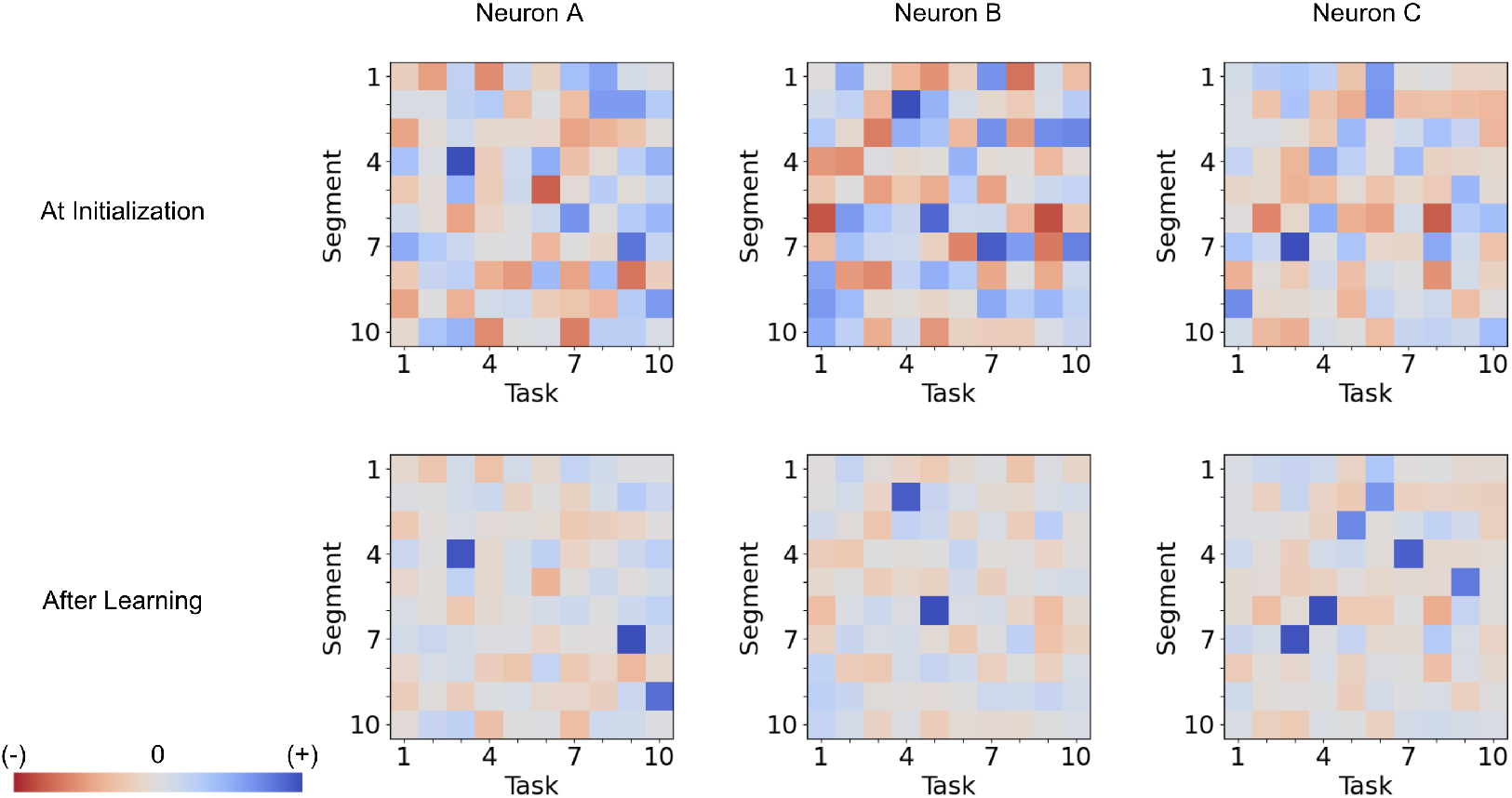
The behavior of the dendritic segments of three separate neurons in the last hidden layer of an Active Dendrites Network for 5,000 randomly-chosen test examples across all tasks, before and after learning 10 permutedMNIST tasks. These charts show the activation computed by each dendritic segment given the context vector corresponding to each task, before (top row) and after (bottom row) training. Note that the dendritic segments for a particular neuron are completely separate of the segments of another (e.g., Neuron A’s first segment is unrelated to Neuron B’s first segment).

### 4.3 Are Networks With Dendrites Equivalent To A Network With More Layers?

Over the last couple of decades, multiple studies have suggested that computations performed by pyramidal neurons can be approximated by ANNs that have one or more hidden layers. Through experiments, Poirazi et al. (2003) showed a two-layer neural network (i.e., multi-layer perceptron (MLP)) can well approximate the pre- and post-synaptic activites of a pyramidal neuron in the hippocampus. Various follow-up studies have also made similar claims (Jadi et al., 2014, Beniaguev et al., 2021). These models suggest that pyramidal neurons have greater capacity and power than a single point neuron. From a computational and deep learning perspective, this is equivalent to claiming that an ANN with dendrites can be substituted by a deeper ANN.

However, we argue that when looking at more realistic dynamic scenarios, such as continual learning, a pyramidal neuron’s activity cannot be approximated by a neural network with multiple layers. Classical deep networks are incapable of performing well in continual learning settings regardless of depth. They are subject to catastrophic forgetting even with more layers. On the contrary, our active dendrites implementation can not only compete with more classical networks in non-continual learning settings, but also perform on-par with state of the art networks on continual learning benchmarks. Figure 9 shows the final accuracy of a network with active dendrites on 10 and 100 permutedMNIST tasks against feedforward networks that have a) the same number of layers but no dendrites, and b) many more layers and roughly the same number of learnable parameters (for 10 tasks).

**Figure 9:**
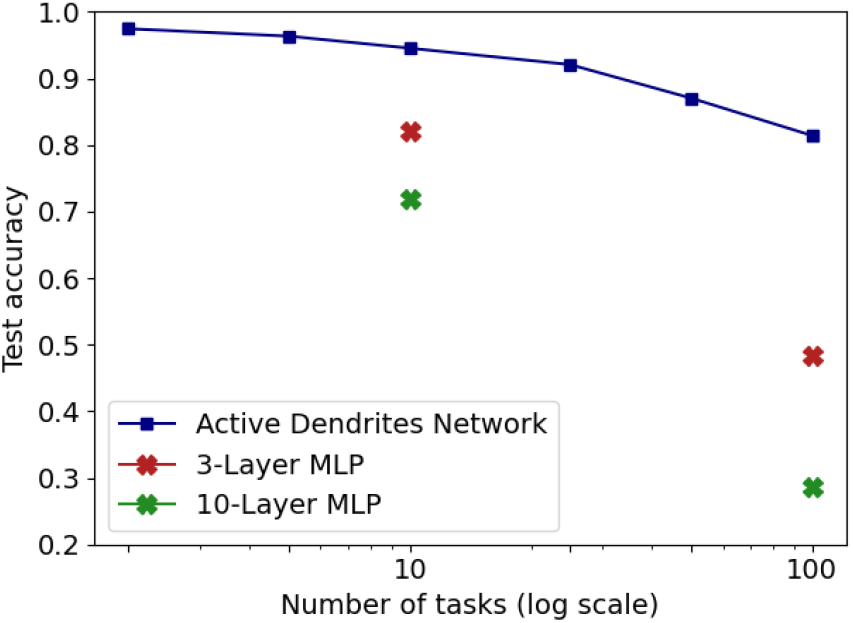
Continual learning overall accuracy for an Active Dendrites Network compared to regular feedforward networks with more layers. Our Active Dendrites Network has three layers. The two hidden layers contain neurons with dendrites.

We also tested other control configurations not shown in Figure 9. First, we tested a 3-layer MLP where either the task ID or context vector was provided as part of the feedforward input. This model performed better than a standard 3-layer MLP but still significantly worse than the Active Dendrites Network. Second, we tested a network with a constant number of learnable parameters independent of the number of tasks (see Appendix C).

Our results suggest that in the realm of continual learning, standard ANNs with multiple layers are prone to catastrophic forgetting while active dendrites can help retain knowledge from previous tasks. Therefore, our Active Dendrites Network is not equivalent a standard feedforward network with more layers.

### 4.4 Comparison With Context Dependent Gating

The idea of leveraging sparse representations and subnetworks within an ANN to combat catastrophic forgetting is not entirely novel. The implementation closest to ours is XdG (Masse et al., 2018) where they hard-coded a distinct subnetwork for each task. When training on a task, they invoked the task-specific subset of the hidden layer of their ANN; other neurons were forced to have an activation value of zero. Their system was provided with a task ID which determined exactly which neurons to turn on or off. Training Active Dendrites Networks in a continual learning scenario also yields subnetworks and sparse representations, however we emphasize two major distinctions between our model and XdG:

1. Task information is inferred in our system (via prototyping) whereas XdG provides the system with a task ID during training and testing. As such, our system is solving a problem that is known to be significantly more challenging (van de Ven and Tolias, 2019).
2. Subnetworks automatically emerge via the use of dendritic segments for each new task whereas XdG pre-allocates a different subnetwork for each task.

We compare Active Dendrites Networks to XdG in Figure 10. Just as we augment Active Dendrites Networks with SI, so too does XdG. Our results with a large number of tasks are significantly better than XdG, and slightly worse than XdG combined with SI, but without their limitations.

**Figure 10:**
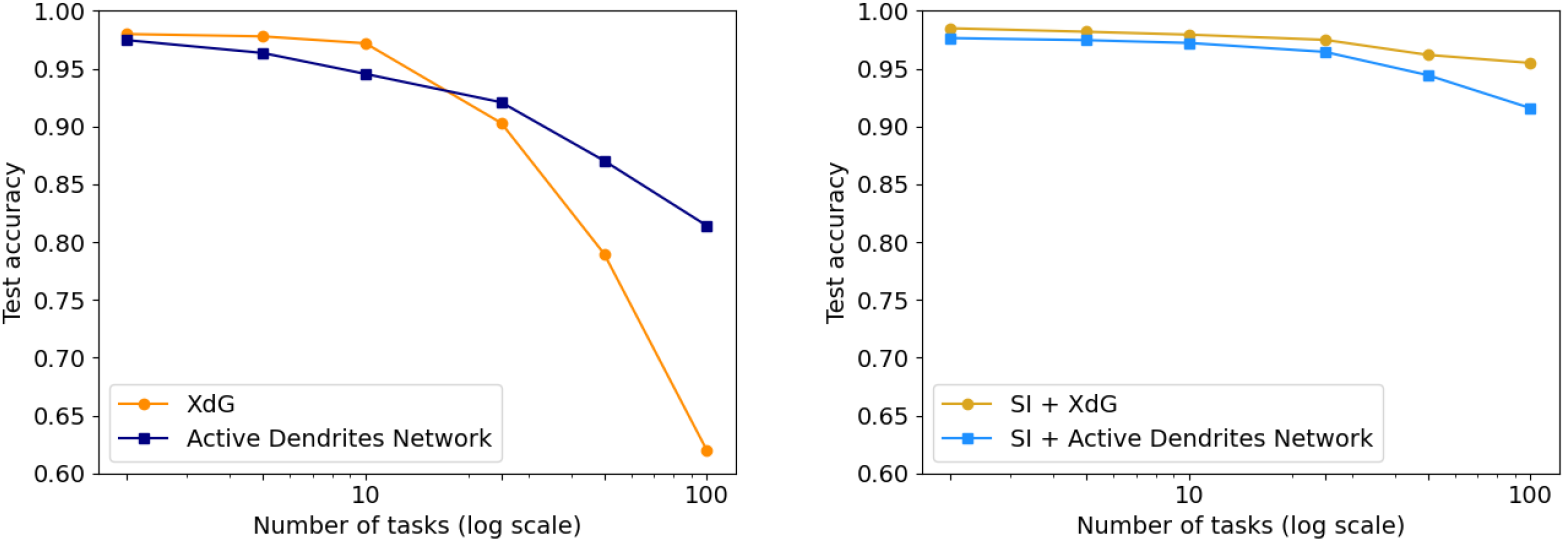
**Left:** Final accuracy of the Active Dendrites Network in comparison to XdG when learning 2, 5, 10, 25, 50, and 100 permutedMNIST tasks. **Right:** Final accuracy of each method when augmented with SI, and SI itself. XdG numbers are taken from Zenke et al. (2017).

Learning is more challenging in our system as dendritic segments must learn the mapping between context vectors and different subnetworks. In effect, sparse representations and minimally overlapping subnetworks emerge naturally in our model. We note that perhaps this makes learning more interesting as dendritic segments can choose subnetworks that overlap more for tasks that are more semantically related, thus requiring less network capacity.

### 4.5 Impact of Sparsity Level and the Number of Dendrites

We have shown that an Active Dendrites Network is competitive with benchmark methods in continual learning. However, to what extent are active dendrites and sparse representations both contributing factors towards alleviating catastrophic forgetting? As it turns out, both 1) active dendrites without sparse representations and 2) sparse representations with standard point neurons are better than chance in a continual learning scenario. However, it is the combination of active dendrites with sparse representations that yield much better results than incorporating just one of these biologically-inspired modeling aspects. Indeed, the accuracy of both methods evaluated independently and combined on up to 100 permutedMNIST tasks clearly demonstrates the importance of having both active dendrites and sparse representations; see Figure 11.

**Figure 11:**
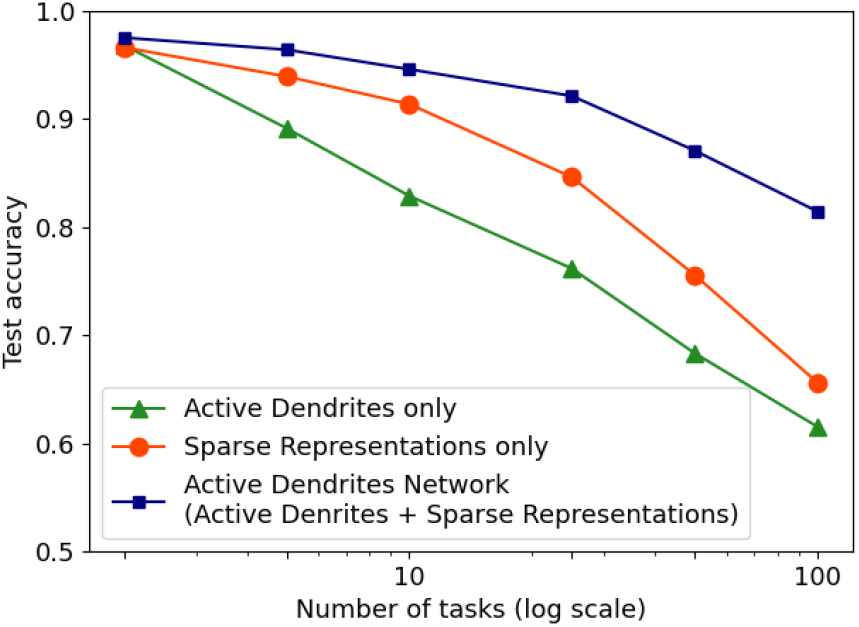
Continual learning mean accuracy on permutedMNIST using active dendrites and dense representations (green), regular ANNs with sparse representations (orange), and Active Dendrites Networks (blue) which use both active dendrites and sparse representations. We average results over 8 independent runs each with a randomly-initialized seed and omit standard error bars as they highlight a very small range.

Furthermore, we also test the effects of varying the number of dendritic segments per hidden neuron (while fixing the level of sparsity in representations), and the result is a small monotonic increase in accuracy. Likewise, decreasing the sparsity level in hidden representations (i.e., increasing *k* in *k*WTA) while keeping the number of dendritic segments constant translates into a sharp drop in accuracy, which highlights the importance of sparse representations. Figure 12 shows results from these experiments when training an Active Dendrites Network on 10 and 50 permutedMNIST tasks in sequence.

**Figure 12:**
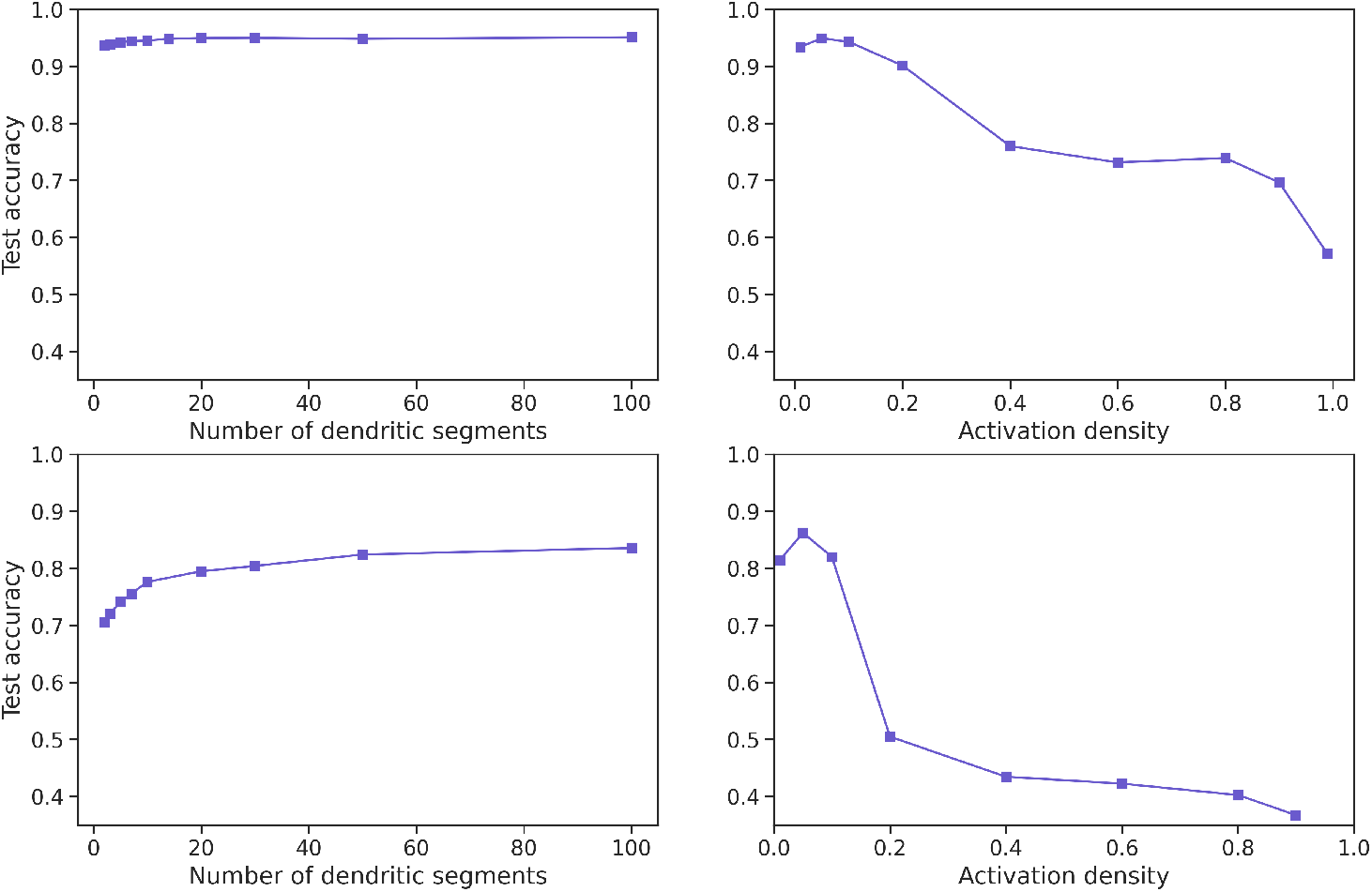
**Left column:** Final accuracy on test examples across all tasks when varying the number of dendritic segments per neuron and keeping activation sparsity constant when learning 10 (top) and 50 (bottom) permutedMNIST tasks. **Right column:** Final accuracy on test examples across all tasks for a fixed number of dendritic segments per neuron and varying activation density level on 10 (top) and 50 (bottom) permutedMNIST tasks.

### 4.6 Understanding Parameters in the Model

One interesting issue relates to the size of the Active Dendrites Network and the total number of parameters. In addition to feedforward weights, our neurons have weights associated with each dendritic segment. In most of our experiments the number of dendritic segments is set to *T*, the number of tasks (See Appendix C for results with a fixed number of dendritic segments). We can calculate the number of weights in each hidden layer *l* of the network as follows. Let *p* be the size of the prototype vector, *n*_*l*_ be the number of units in layer *l, s*^*F*^ be the weight sparsity for the feedforward weights and *s*^*D*^ the weight sparsity for dendritic weights. The total number of weights in layer *l* is then:

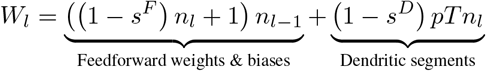

The first term represents the total number of weights in the feedforward portion (including a bias). The second term represents the number of weights in the dendritic segments. In our implementation, *s*^*F*^ = 0.5, and *s*^*D*^ = 0 (i.e., dendritic weights are fully dense).

In addition to these weights we also store *T* prototypes, each of which has the same size as the input vector. Although these are not learned through backpropagation, they are determined from the training data and should be included in the parameter count. In permutedMNIST the input vector size is *n*_0_ = 784, leading to a total of *T ×* 784 additional values for the prototypes.

The number of dendritic weights quickly dominates all other parameters as the number of tasks increases (Table 1, middle column). At first glance, the implication is that the number of parameters in our 100-task network is far greater than the number of parameters in the comparison networks. However notice that the dendritic segments do not receive the input. The dendritic segments determine a context-dependent scale factor per neuron, based only on one of 100 possible context vectors. This scale factor is learned during training but then is static during testing.

**Table 1:** The total and effective number of parameters for Active Dendrites Networks as compared to some of the other networks. Note: that XdG requires a mapping from task ID to sub-networks that is not incorporated in the table.

| Network | Tasks | Parameters | Effective Parameters |
| --- | --- | --- | --- |
| Active Dendrites Network | 10 | 35,034,794 | 2,963,114 |
| Active Dendrites Network | 100 | 324,119,114 | 3,402,314 |
| 3-layer MLP | 10 or 100 | 5,824,512 | 5,824,512 |
| 10-layer MLP | 10 or 100 | 35,198,976 | 35,198,976 |
| XdG | 10 or 100 | 5,592,010 | 5,592,010 |

Since there is a small fixed pool of *T* prototype vectors, a simple post-processing step can replace the weights with a smaller identical system. From Section 3.1, the output of a neuron is:

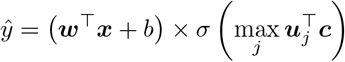

During testing 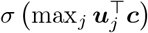 is constant for each vector ***c***_*i*_. The equation can be re-written as:

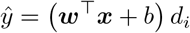

where 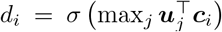, 0 < *i* ≤ *T*. For any given test input, we can select the nearest prototype vector *i* and use the appropriate scale factor *d*_*i*_. The total number of effective parameters in layer *l* is thus reduced to:

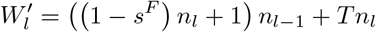

Note that in the experiments reported here, with a small fixed number of context vectors, it is actually possible to learn *d*_*i*_ directly via backpropagation. In this case the number of parameters would be identical to the number of effective parameters, even during training. We did not implement this as it would also limit the flexibility of the overall architecture and disallow future scenarios where the context vector changes dynamically per input. In Table 1 we list the total and effective number of parameters for the Active Dendrites Network in comparison to some of the other networks. Note that the Active Dendrites Network has substantially fewer effective parameters than any of the other networks.

In our previous work (Hawkins and Ahmad, 2016) we used extremely sparse dendritic weights (*>* 99% sparsity). These weights were dynamically determined during the learning process by sampling from components of the context vector. Consistent with the biology of active dendrites, the number of weights per segment was limited to a small constant (such as 30). Implementing sparse dendritic weights in the context of deep learning systems is an important future research area for Active Dendrites Networks.

## 5. Discussion

Ever since Rosenblatt (Rosenblatt, 1958), AI visionaries have attempted to build intelligent systems by taking ideas from neuroscience and turning them into algorithms. They looked to neurons in the brain as a single computational unit and constructed ANNs by organizing multiple such units in a hierarchy. The exact mechanistic details of how a biological neuron converts incoming signals into action potentials (i.e., spikes) have always been unclear. Implementations of biological neurons in silico have favored a single linear weighted sum (the point neuron) as a tractable abstraction. This idea continues to serve as the prevalent paradigm in machine learning today for the individual computational unit.

One shortcoming is that standard ANNs with point neurons overwrite most of their connections for each learning iteration, and thus quickly lose previously-acquired knowledge (French, 1999, Parisi et al., 2019). In this paper we have shown that augmenting point neurons with biological properties such as active dendrites, complex synapses, and sparse representations significantly improves a network’s ability to learn continually. In particular, we have shown that a 3-layer Active Dendrites Network with SI can achieve greater than 90% accuracy when learning 100 permutedMNIST tasks in sequence. In the following subsections we discuss some intuitions for why dendrites help with catastrophic forgetting, and some relationships to other papers.

### 5.1 Dendrites Invoke Subnetworks

In this section we attempt to shed light on why active dendrites help in continual learning. In our model dendritic segments in each neuron identify specific contexts and then modulate neuronal activity based on this identification. Due to the subsequent *k*-Winner-Take-All function, the modulation can have an impact on whether the neuron will become active. We propose that the impact of this behavior is to invoke sparse context-specific subsets of the network. This in turn causes learning to be highly localized and task-specific, and helps reduce forgetting effects.

Figure 13 visually illustrates these subnetworks. Two different context vectors can lead to different winners and different sparse activation patterns. As suggested by the figure, it is even possible for the exact same feedforward input to activate completely different neurons for different contexts. Note that the subnetworks are distributed; two different subnetworks may have neurons in common.

**Figure 13:**
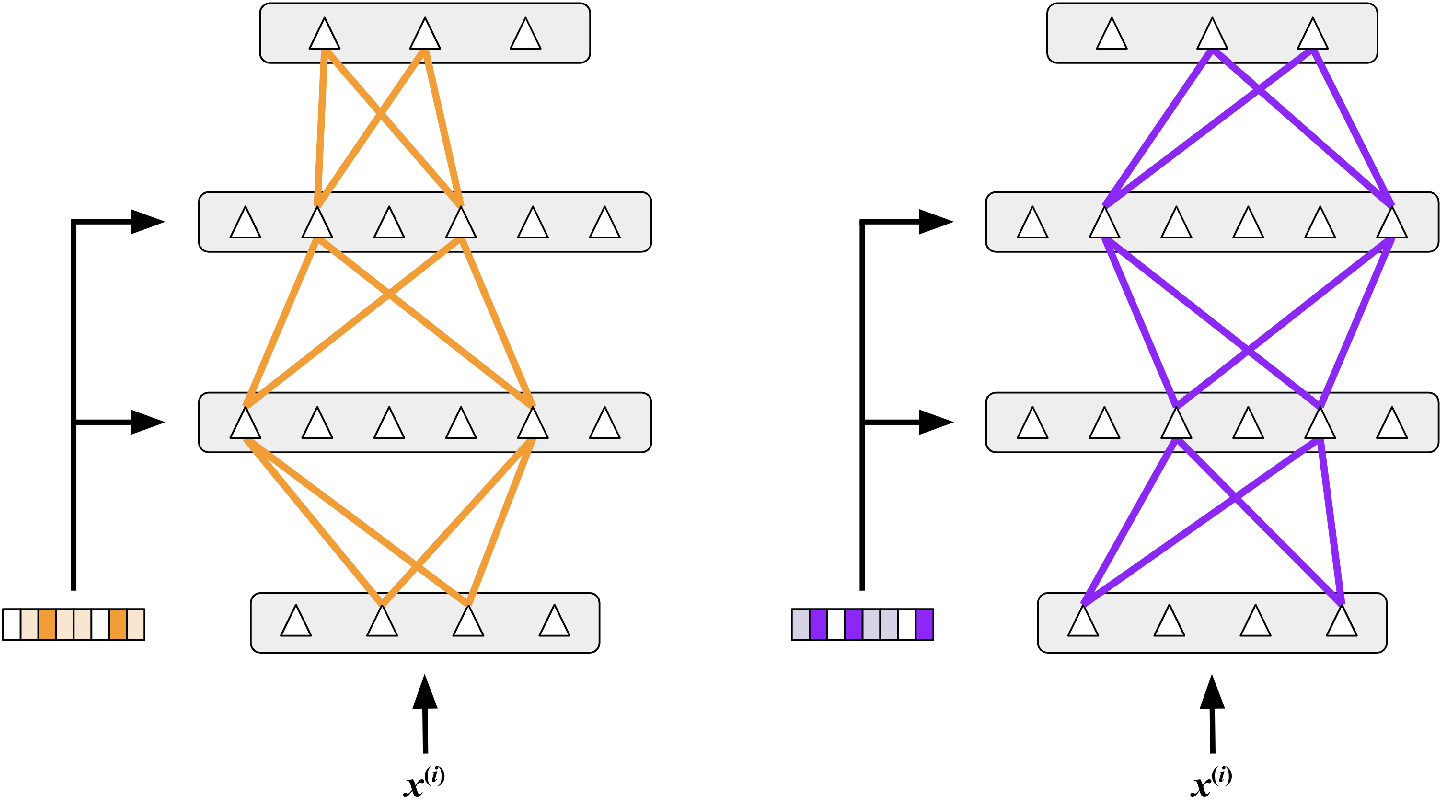
A cartoon illustration of subnetworks within an Active Dendrites Network. By receiving different context vectors as input, dendritic segments can invoke different subnetworks for a fixed feedforward input. The subnetworks are distributed, i.e., they may share some of the same neurons.

In Section 4.2 we showed that task-specific representations do indeed emerge (Figure 6), and that the representations from these subnetworks have minimal overlap (Figure 7). Our analysis also showed that once learning is complete, individual segments within a neuron respond strongly to different context vectors (Figure 8).

What is remarkable is that even though dendritic segments receive a particular context vector only when training on the corresponding task, they can still retain that information much later in the process, and correctly activate the appropriate subnetwork. We believe the reason for this is that*·* the *k*WTA function prevents the majority of the neurons from activating, thus error will only be backpropagated through the active subnetwork. In addition the max(·) operator causes errors to be backpropagated only to the winning dendritic segment. Thus, only the active neurons will update their feedforward weights. Only the winning segment within those active neurons will update their dendritic weights. We suspect that this highly selective weight update helps mitigate the effect of catastrophic forgetting. Interestingly, there is experimental support that this type of highly localized learning occurs in pyramidal neurons (Losonczy et al., 2008, Limbacher and Legenstein, 2020).

### 5.2 Comparing a Neuron With Active Dendrites to ANNs

Previous studies have proposed that a single pyramidal neuron with active dendrites is computationally equivalent to an ANN with multiple layers (Mel, 1992, Poirazi et al., 2003, Poirazi and Papoutsi, 2020, Jones and Kording, 2021). They suggest that the non-linear integrative properties in dendritic segments add depth and increase capacity, thereby allowing a neuron to implement more complex input-output functions than a single point neuron. Interestingly, from a practical machine learning perspective this implies that active dendrites are not fundamentally different. It is sufficient to simply substitute dendritic neurons with a deeper network of point neurons. In part due to this result, dendrites have not become popular in deep learning.

In this paper we propose that active dendrites can in fact add value, particularly in more realistic dynamic settings. In particular, the equivalence to deeper networks does not hold in continual learning. We showed in Section 4.3 that Active Dendrites Networks significantly outperform regular ANNs with more hidden layers when learning multiple tasks in sequence. It Section and 4.6 we showed that this is true even when the effective capacity of the Active Dendrites Network is significantly smaller than a standard ANN. Solving the catastrophic forgetting problem requires added functionality that is independent from increasing capacity. We suggest that the subnetwork functionality described above goes beyond simply adding layers and parameters. Instead they may play a critical role in the brain in handling complex dynamic scenarios.

### 5.3 Related ANN Architectures

Another way to think about neural networks with Active Dendrites Neurons is that they dynamically decide on a representation for the feedforward inputs based on context. When the modulation function *f* involves multiplication, dendritic networks fall under the umbrella of multiplicative networks. Jayakumar et al. (2020) demonstrated that multiplicative networks can excel in multitask learning by leveraging dynamic representations in a task-specific manner. Two prominent examples of dynamic networks include FiLM layers (Perez et al., 2018) and Transformers (Vaswani et al., 2017). This framing also suggests further investigating the overlap between multitask learners that receives side information about the task at hand and multimodal learners that fuse multiple streams of input from different sensory modalities. Conversely, the dendritic networks explored here could be explored in the context of multimodal learning, or other topic areas where dynamic networks have been successful.

A related technique which has also been widely used in continual learning is gating. Many works in this vein draw on the mixture of experts framing (Jacobs et al., 1991). Gated Linear Networks (Veness et al., 2021) and Dendritic Gated Networks (Sezener et al., 2021) are noteworthy examples of networks that use context-specific gating as these models are not trained through standard back-propagation. Our architecture is related in that the dendritic ouput gates the activation of a neuron. One notable difference is that the neuronal activations are sparse in our model.

In our case, dendritic networks create input representations composed of entirely different subnetworks of neurons. These sparse representations are a special type of dynamic representation. Supermasks and XdG also explicitly utilize sparse subnetworks per task. XdG hard-codes subnetworks for each task, and this extra supervision step forgoes the need to dynamically gate activations and makes training/testing slightly easier. Supermasks also maintains an explicit list of masks to use outside the network. By contrast, our model invokes sparse subnetworks automatically and without an explicit.

### 5.4 Future Work

Although we have shown an initial proof of concept that active dendrites and sparse representations can help in continual learning, there is still significant work to be done. First, we only test our model empirically on permutedMNIST. Despite being a benchmark continual learning dataset, it is far from the types of inputs a robust learner should be able to perform well on, such as real world images. Second, our prototype method for computing a context signal is simple and works well in practice, but may not scale to more challenging problems. We suspect that the simplicity of permutedMNIST images provides good task separation and this leads to the prototype method being successful in the scenarios we trained in, but a more principled approach towards constructing context vectors for a broad range of datasets remains to be investigated. Finally, we discussed the role of sparse representations throughout this paper, but have not explored sparse weights in depth (Ahmad and Scheinkman, 2019). In our experiments, feedforward weights were 50% sparse, however dendritic segments were entirely dense. In future work, we would like to address how to create sparse dendritic segments.

## A Absolute Max Gating

Here, we briefly outline how we implement gating in Active Dendrites Networks. In section 3, we originally present gating as modifying the value of the weighted linear sum computed by the point neuron based on the maximum activation, i.e., σ(max_*j*_ ***u*** ^⊤^ ***c***). However, one clear problem with this formulation is that it becomes difficult to turn a neuron off (i.e., force it’s activation value to be zero) due to the max operator. That is, if dendritic segment *j* learns to turn off the unit, then based on sigmoidal gating, we should expect that 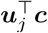 is a small number with large absolute value (very negative). However, there’s a very good chance that for some segment *j*^*′*^ (where *j ≠ j*^*′*^), 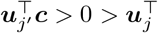 which not entirely turn the neuron off and increase the chance it becomes selected by the *k*WTA process.

This motives absolute max gating in which the activation with the largest magnitude is selected and its sign is kept. More formally, a point neuron augmented with absolute max gating comptutes its output as

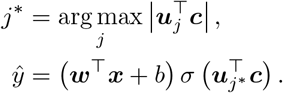

## B Inferring Prototypes during Training

When task information is not given during training nor testing, the task corresponding to each input example must be inferred. Section 3.3 describes how this was done at test time, and this section describes an online method to estimate task information during training. One inductive bias in our procedure is that all training examples in a batch correspond to the same task, since continual learning scenarios usually only observe examples from a single task within a given batch.

### B.1 Clustering Approach

Formally, let *X* = *{****x***^(1)^, …, ***x***^(*n*)^*}* be a batch of *n* training examples (in the case of permutedMNIST, each ***x***^(*i*)^ is a 784-dimensional vector for 1 *≤ i ≤ n*). Suppose *M* individual prototypes have been designated thus far: ***p***_1_, …, ***p***_*M*_. For each ***p***_*j*_ (where 1 *≤ j ≤ M*), the individual examples used to construct that prototype are also stored in memory: 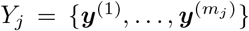, where *m*_*j*_ gives the number of examples for cluster *j*. These previous training examples were observed by the learner during previous batches of learning and have been stored in memory. We want to identify if the new batch *X* is similar enough to any cluster of training examples *Y*_*j*_ such that the corresponding prototype ***p***_*j*_ should be used as the context signal. If a cluster *j* is found such that *X* is “similar” to *Y*_*j*_, then *Y*_*j*_ is expanded to include *X*. Subsequently, ***p***_*j*_ is updated to incorporate samples from *X*. Otherwise, if *X* is deemed significantly different from *Y*_*j*_ for all *j*, then a new cluster is formed: *Y*_*M*+1_ *←X* and its prototype is is the element-wise mean of all ***x*** *∈ X*. Algorithm 1 describes the procedure for clustering during training when task information is not provided.

#### Algorithm 1

Clustering algorithm by which a new batch of inputs *X* either gets assigned to 1 of *M* existing clusters or initiates cluster *M* + 1. This procedure is greedy since it assigns *X* to the first cluster *j* that it suitably matches.

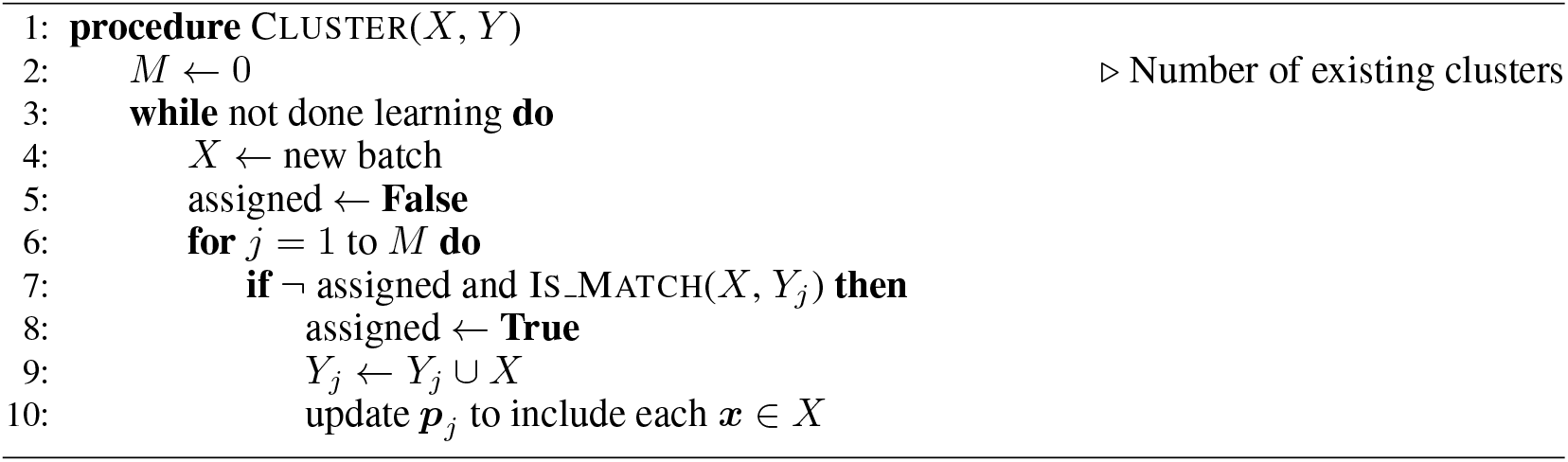

### B.2 Multivariate t-Test

In the above pseudocode, how do we determine when *X* is similar enough to some *Y*_*j*_? If we had univariate data (i.e., if each ***x*** ∈ *X* and ***y*** ∈ *Y*_*j*_ was a scalar quantity), we could use an unpaired *t*-test do this. Instead, we use a generalized version of an unpaired *t*-test that applies to multivariate data.

In our hypothesis testing setup, the null hypothesis is that for any given *j*, the same underlying process generates samples from both *X* and *Y*_*j*_. When we accept the null hypothesis, we assume each ***x*** *∈ X* and each ***y*** *∈ Y*_*j*_ are training examples from the same permutedMNIST task—and therefore ***p***_*j*_ can be used as the context signal when training an Active Dendrites Network on examples in *X* (albeit ***p***_*j*_ is first updated to account for *X*).

Hotelling (1931) proposed Hotelling’s *t*-squared statistic (*t*^2^) as a generalization of the *t*-statistic used to perform single-variable *t*-tests; it is computed as

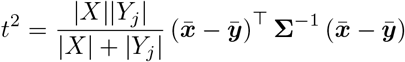

where 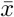 and 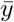 are simply the element-wise means of all ***x*** *∈ X* and ***y*** *∈ Y*_*j*_, respectively, and **Σ** is the pooled, sample-adjusted covariance matrix of samples in *X* and *Y*_*j*_. The test statistic *t*^2^ can be compared to a chosen *p*-value to accept or reject the null hypothesis by first transforming it to a value drawn from an *F* -distribution (whose cumulative density function is more well-studied than that of the *t*-squared distribution) as follows:

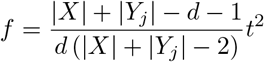

where *d* is the dimensionality of the samples.

We fix a *p*-value and derive a value for *f* based on *t*^2^ as give above. If *f > p*, then we reject the null hypothesis since the probability that the same generative process explains both *X* and *Y*_*j*_ is extremely low, and thus create a new cluster. Since we perform pairwise multivariate *t*-tests between *X* and *Y*_*j*_ for all existing prototypes *j*, a new cluster and prototype emerge if and only if we reject the null hypothesis for all *M t*-tests. Algorithm 2 describes the procedure for performing the multivariate *t*-test via the *t*-squared statistic given two sets of multivariate samples.

#### Algorithm 2

Unpaired multivariate *t*-test using Hotelling’s *t*-squared statistic. Here, we use a slight abuse of notation when computing covariance matrices by assuming sets of *d*-dimensional vectors can also be treated as matrices whose rows correspond to their *d*-dimensional elements. We assume a *p*-value is fixed a priori.

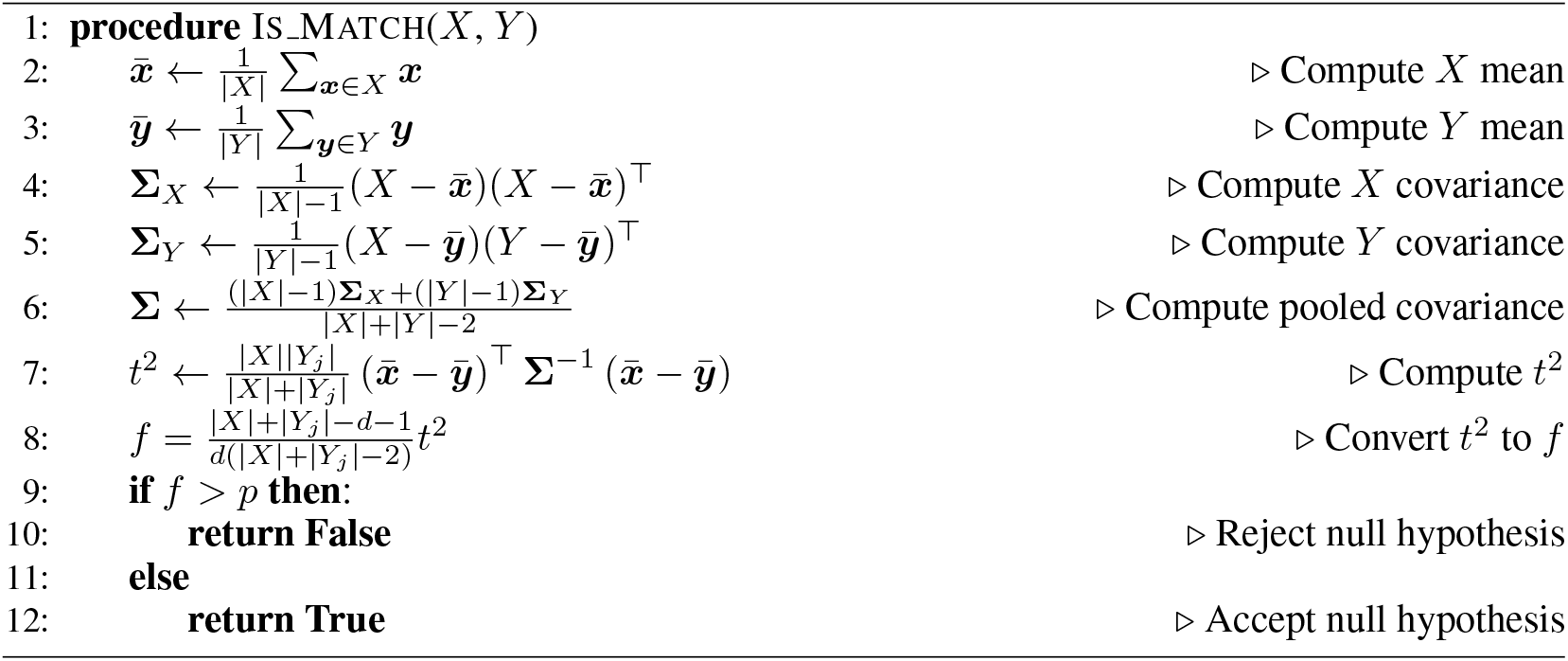

We remark that in our implementation, we replace all standard matrix inversions with the Moore-Penrose pseudo-inversion.

### B.3 Number of Clusters Formed

When employing Algorithms 1 and 2 to infer the context vector while training, we used a significance threshold of *p* = 0.9. We chose this value arbitrarily, and can further improve our results by incorporating *p* as a model hyperparameter. Assuming that the prototype vector for each permutedMNIST task is sufficiently different, we found that our method arrives at a “sensible” number of prototypes (i.e., not too few nor too many clusters/prototypes as compared with the number of tasks). Figure 14 shows the average number of clusters formed as a function of the number of permutedMNIST tasks we trained an Active Dendrites Network on.

**Figure 14:**
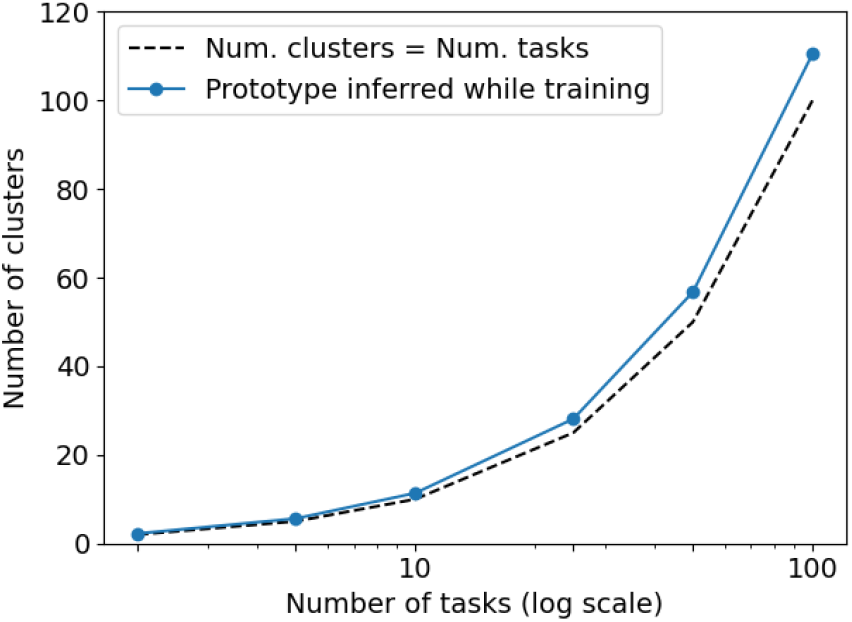
The average number of clusters found by the clustering procedure (described by Algorithms 1 and 2) as a function of the number of permutedMNIST tasks. All results are averaged over 8 independent trials.

## C Active Dendrites Network with a Fixed Number of Parameters

In our experiments we mentioned that for any given number of permutedMNIST tasks, a single Active Dendrites Neuron has the same number of dendritic segments as tasks. The total number of learnable parameters in that scenario grows linearly with the number of tasks. (Table 1 lists each model’s parameter count.) Although the number of effective parameters is far smaller than the actual parameter count (see Section 4.6), we also tested learning 100 tasks in sequence with a fixed 10 dendritic segments per neuron. This network maintains a constant 35 million parameters (same as a 10-layer MLP) independent of the number of tasks. As Figure 15 shows, our modified network achieves 78.5% accuracy on 100 tasks, close to the network with 100 dendritic segments.

**Figure 15:**
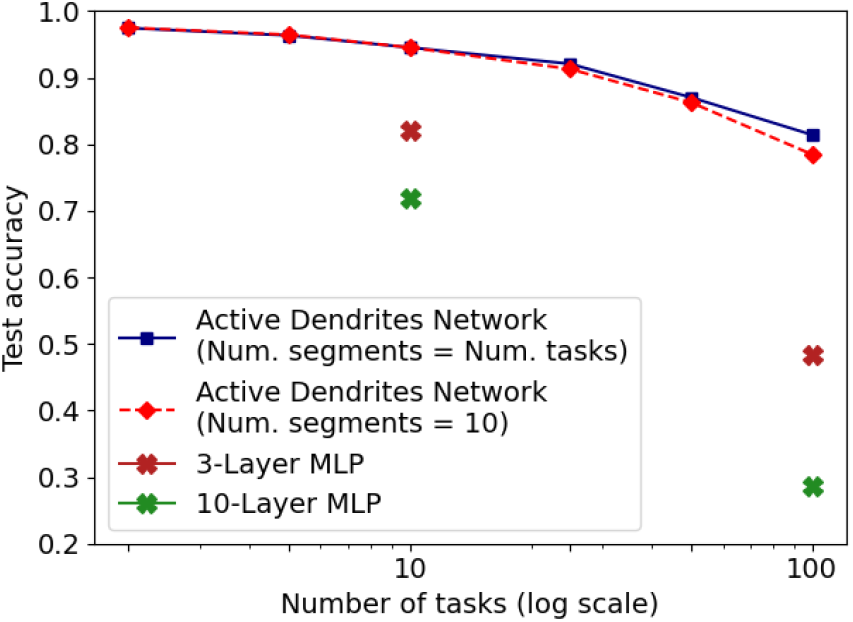
Continual learning accuracy on permutedMNIST: an Active Dendrites Network with the same number of dendritic segments per neuron as the number of tasks (blue), and one with exactly 10 dendritic segments per neuron (red). We also included 3- and 10-layer MLPs on 10 and 100 tasks. All results are averaged over 8 independent trials.

Why does an Active Dendrites Network not suffer from a severe drop in accuracy with significantly fewer dendritic segments for a large number of tasks? We hypothesize that since the dendritic segments are dense and prototype context vectors are sparse (as most pixels in an MNIST image are black), a single segment can learn to identify multiple context vectors, and thus there can be far fewer dendritic segments than unique context vectors.

## D Hyperparameters

### D.1 Active Dendrites Networks

In this section, we provide hyperparameters that we use to train Active Dendrites Networks. When employing the prototype method to infer a context signal at test time only, we train a three-layer network with 2,048 units in each of the two hidden layers, and 10 units in the output layer. The context signal was a 784-dimensional vector, and each training batch consists of 256 examples. After each of the two hidden layers, we apply *k*WTA as our choice of non-linear function with *k* = 102. It’s worth noting that despite the number of permutedMNIST tasks to learn in sequence, we use a single output head and do not freeze any weights, as is done frequently in many continual learning scenarios. Feedforward weights were 50% sparse (i.e., only half were non-zero) and dendritic segments were entirely dense. The number of dendritic segments per hidden unit was set equal to the number of tasks to learn in sequence. We use the Adam optimizer (Kingma and Ba, 2015) and the table below provides the learning rate and number of training epochs per task.

| Num. Tasks | 2 | 5 | 10 | 25 | 50 | 100 |
| --- | --- | --- | --- | --- | --- | --- |
| Learning Rate | $5 \times 10^{-4}$ | $5 \times 10^{-4}$ | $5 \times 10^{-4}$ | $3 \times 10^{-4}$ | $3 \times 10^{-4}$ | $10^{-4}$ |
| Num. Epochs | 1 | 1 | 3 | 5 | 3 | 3 |

In Appendix C we present results with Active Dendrites Networks that use exactly 10 dendritic segments per neuron, and this next table gives the corresponding hyperparameters.

| Num. Tasks | 2 | 5 | 10 | 25 | 50 | 100 |
| --- | --- | --- | --- | --- | --- | --- |
| Learning Rate | $10^{-3}$ | $7 \times 10^{-4}$ | $5 \times 10^{-4}$ | $3 \times 10^{-4}$ | $7 \times 10^{-5}$ | $7 \times 10^{-5}$ |
| Num. Epochs | 5 | 5 | 3 | 5 | 5 | 3 |

When task information is not provided during training nor test time and we employ the clustering procedure from Appendix B to construct prototypes during training, we again use the same architecture, optimizer, and batch size as before but with the learning rates and training epochs per task listed in the table below. Also, we decrease the the number of context dimensions to 256 by randomly sampling features from the original 784-dimensional input. The reason for this decrease is that when converting a *t*^2^ value to an *F* -distributed variable as in Algorithm 2, the context dimension must be less than the the sum of sizes of two sets (512 in our case).

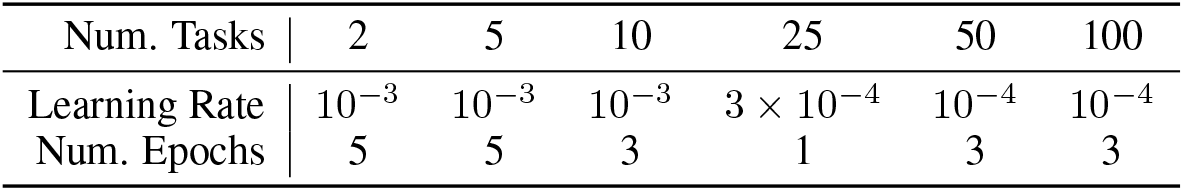

To combine Active Dendrites Network with SI, we reduce the number of units in each hidden layer from 2,048 to 2,000 as to exactly match the architectures (with the exception of dendritic segments) of the network used in the SI and XdG experiments. In addition, the SI-and-Active-Dendrites network was trained for 20 epochs per task instead of just 3 (with Active Dendrites Networks) as this significantly improved results. We fix the learning rate to be 5 *×* 10^*−*4^ for all numbers of tasks, and we use SI regularization strength *c* = 0.1 and damping coefficient *ξ* = 0.1. Both a) training for 20 epochs per task and b) the *c, ξ* values that we use here align with the training setups of Zenke et al. (2017) and Masse et al. (2018).

### D.2 MLPs

Section 4.3 describes comparing continual learning results of a network with active dendrites vs one with more layers. In these experiments, we train a MLPs with 3 layers and one with 10 layers, each with 2,048 hidden units in each hidden layer and 10 units in the output layer. These MLPs had fully dense weights (unlike Active Dendrites Networks) and and used a ReLU non-linear activation function in each hidden layer. To find a suitable choice of hyperparameters to train these MLPs, we perform a 2D grid search over the number of training epochs per task (up to 25 epochs) and the learning rate (in the range [10^*−*7^, 10^*−*3^]). Just as with Active Dendrites Networks, we train MLPs with Adam, and the following table lists the learning rate and number of training epochs per task used to obtain accuracy in Figure 9.

| Num. Tasks |  | 10 | 100 |
| --- | --- | --- | --- |
| 3-Layer MLP | Learning Rate | $3 \times 10^{-6}$ | $10^{-6}$ |
|  | Num. Epochs | 5 | 3 |
| 10-Layer MLP | Learning Rate | $3 \times 10^{-6}$ | $3 \times 10^{-7}$ |
|  | Num. Epochs | 3 | 3 |

### D.3 Ablation Studies

Figure 11 shows results from an ablation study in which we compare Active Dendrites Networks to both a network that has uses active dendrites for gating but no sparse representations (AD) and a regular feedforward network that uses only sparse representations (SR). When training these networks in the same continual learning scenario as the Active Dendrites Network, we use the same network architecture as before, with the exception that the SR model had no dendritic segments. The table below gives learning rates and number of training epochs per task for SR and AD models.

| Num. Tasks |  | 2 | 5 | 10 | 25 | 50 | 100 |
| --- | --- | --- | --- | --- | --- | --- | --- |
| Learning Rate | AD | $5 \times 10^{-4}$ | $10^{-3}$ | $5 \times 10^{-6}$ | $5 \times 10^{-6}$ | $10^{-5}$ | $5 \times 10^{-6}$ |
| | SR | $10^{-3}$ | $5 \times 10^{-5}$ | $10^{-4}$ | $10^{-5}$ | $10^{-5}$ | $10^{-5}$ |
| Num. Epochs | AD | 5 | 10 | 5 | 5 | 5 | 5 |
|  | SR | 3 | 10 | 5 | 10 | 10 | 5 |

## Footnotes

1 GitHub Repository: https://github.com/numenta/htmpapers

